# *ATRX, DAXX or MEN1* mutant pancreatic neuroendocrine tumors are a distinct alpha-cell signature subgroup

**DOI:** 10.1101/195214

**Authors:** Chang S Chan, Saurabh V Laddha, Peter Lewis, Matthew Koletsky, Kenneth Robzyk, Edaise Da Silva, Paula J. Torres, Brian Untch, Promita Bose, Timothy A. Chan, David S. Klimstra, C. David Allis, Laura H. Tang

## Abstract

The most commonly mutated genes in pancreatic neuroendocrine tumors (PanNETs) are *ATRX*, *DAXX*, and *MEN1*. Little is known about the cells-of-origin for non-functional neuroendocrine tumors. Here, we genotyped 64 PanNETs for mutations in *ATRX*, *DAXX*, and *MEN1* and found 37 tumors (58%) carry mutations in these three genes (A-D-M mutant PanNETs) and this correlates with a worse clinical outcome than tumors carrying the wild-type alleles of all three genes (A-D-M WT PanNETs). We performed RNA sequencing and DNA-methylation analysis on 33 randomly selected cases to reveal two distinct subgroups with one group consisting entirely of A-D-M mutant PanNETs. Two biomarkers differentiating A-D-M mutant from A-D-M WT PanNETs were high *ARX* gene expression and low *PDX1* gene expression with *PDX1* promoter hyper-methylation in the A-D-M mutant PanNETs. Moreover, A-D-M mutant PanNETs had a gene expression signature related to that of alpha cells (pval < 0.009) of pancreatic islets including increased expression of HNF1A and its transcriptional target genes. This gene expression profile suggests that A-D-M mutant PanNETs originate from or transdifferentiate into a distinct cell type similar to alpha cells.

## Introduction

Pancreatic neuroendocrine tumors (PanNETs) or Islet Cell Tumors are a rare neuroendocrine malignancy with an annual incidence of less than 1 per 100,000 per year^1^ (about 1,000 new cases per year in the United States → 3 cases per million.) Current classification scheme for PanNETs include grade and stage^1^. While well-differentiated PanNETs can be successfully treated with surgery, there are few treatments for metastatic PanNETs. A greater understanding of PanNET pathogenesis may guide the development of novel therapeutic options.

Molecular studies have identified mutations in *MEN1*, *ATRX*, and *DAXX* to be the most commonly found in PanNETs^2^,^3^ (found in approximately 10, 20 and 40% of tumors, respectively). All three genes play a role in chromatin remodeling. MEN1 is a component of a histone methyltransferase complex^4^ that specifically methylate Lysine 4 of histone H3 and functions as a transcriptional regulator. ATRX and DAXX interact to deposit histone H3.3-containing nucleosomes in centromeric and telomeric regions of the genome^5^. Additional mutations in mTOR pathway genes including *TSC2, PTEN,* and *PIK3CA* are found in one in six well-differentiated PanNETs^2^.

Most PanNETs are classified as nonfunctional as they do not cause any symptoms until later stages of tumorigenesis, while a minority cause symptoms due to excessive secretion of hormones and are thus functional. The neuroendocrine cells in the pancreas include alpha, beta, delta, pancreatic polypeptide (pp)-producing and vasoactive intestinal peptide (VIP)-producing cells. The cell of origin for nonfunctional PanNETs is not well established. Here, we genotyped 64 well differentiated PanNETs for mutations in *ATRX*, *DAXX* and *MEN1* and performed RNA sequencing (n=33) and DNA methylation (n=32) analysis to identify distinct molecular phenotypes of A-D-M mutant PanNETs which potentially reveals their distinct cell of origin or transdifferentiated state.

## Results

### Patient cohort, clinical annotation, and genotyping for ***ATRX***, ***DAXX* and *MEN1***

We initially performed targeted sequencing to genotype the *ATRX*, *DAXX* and *MEN1* genes in 64 individual PanNETs. All cases were histologically confirmed to be well-differentiated PanNETs of WHO G1/G2 grade, and cases of poorly differentiated neuroendocrine carcinoma were excluded. The mean patient age was 52±1.5 years (ranging from 26-73) with a 59% male population. The locations of the tumors were 38% proximal/mid body, and 62% distal pancreas. Eighty-one percent of the cases were clinically non-functional and the remaining cases included insulinomas, glucagonomas, gastrinomas, and VIPomas. The median size of tumor was 3.6±0.4 cm (ranging from 1.0 – 14.5 cm). Sixty-eight percent of patients had localized disease without distant metastasis at the time of initial diagnosis (Supplementary file 1).

An A-D-M mutant genotype was identified in 58% (37/64) of cases with *ATRX, DAXX, MEN1, MEN1/ATRX,* and *MEN1/DAXX* mutations in 8%, 16%, 20%, 3%, and 11% cases, respectively (Figure 1a). Among 44 patients who initially presented with localized disease without distant metastasis, those with the A-D-M mutant genotype had a worse recurrence free survival than those of A-D-M WT genotype (Figure 1b).

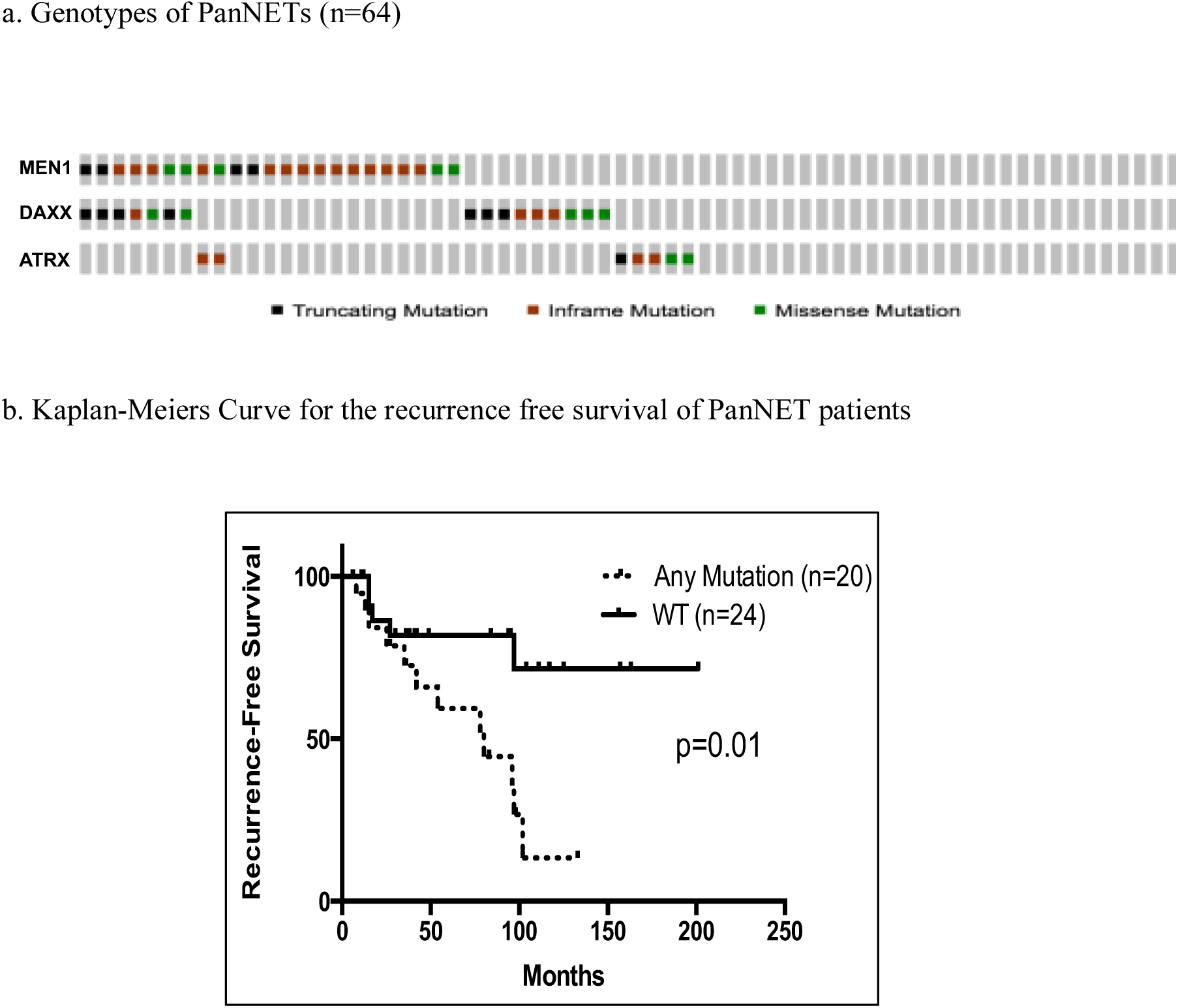

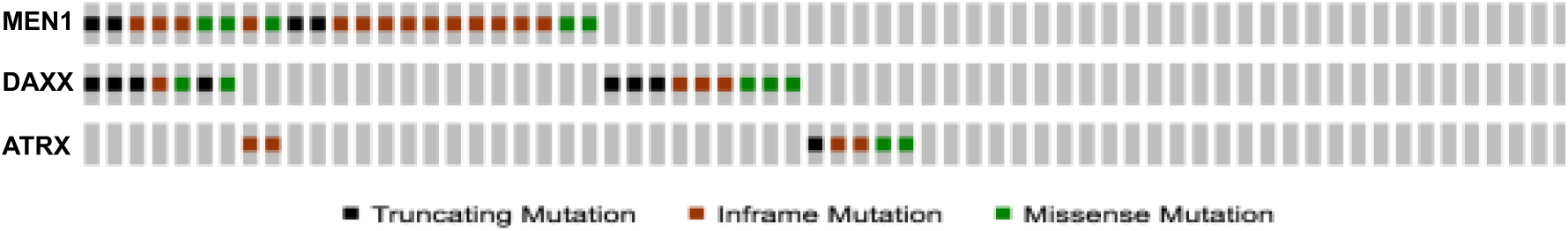

**Figure 1.**
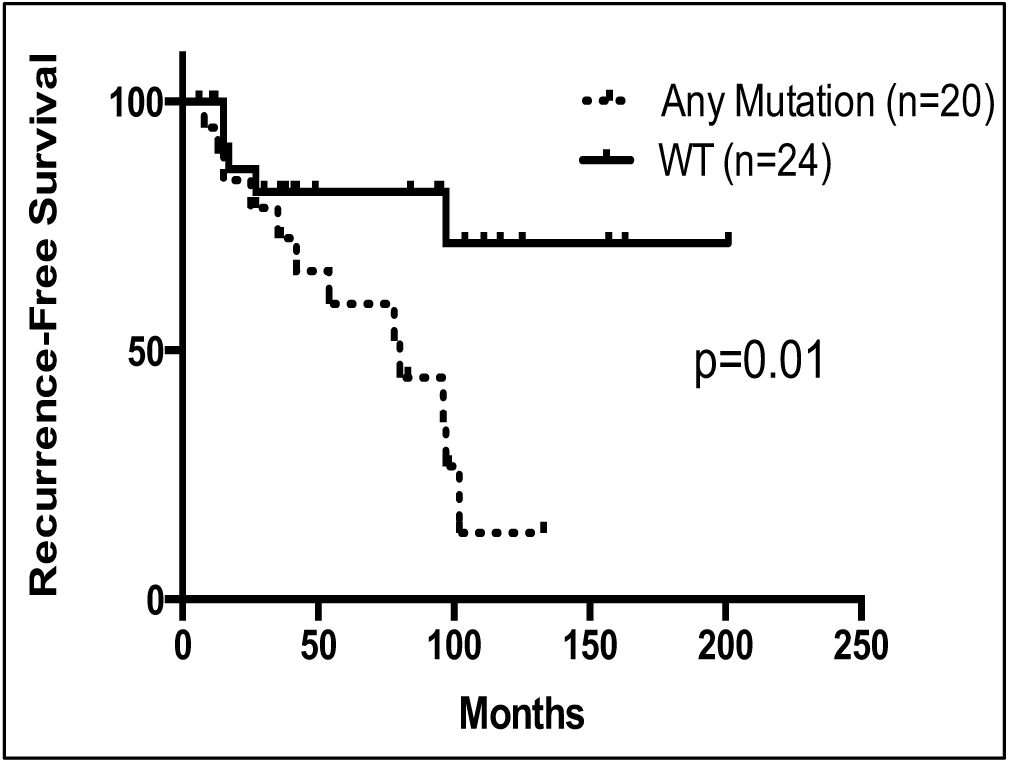
Mutational landscape of ATRX, DAXX and MEN1 in PanNETs. a) Oncoprint mutational profile for PanNETs samples. *ATRX/DAXX/MEN1* mutations were identified in 37/64 (58 %) of PanNETs using Sanger and miSeq sequencing. b) Among 44 patients who initially presented with localized PanNETs (without distant metastasis), those with A-D-M mutant genotype had a worse recurrence free survival outcome than those A-D-M wt genotype in their primary tumors. A-D-M mutated samples are annotated as any mutation (n=20) and A-D-M WT samples annotated as WT (n=24).

### Gene expression and DNA methylation reveal two subtypes of PanNETs

We performed RNA sequencing on 33 tumors by random selection (19 A-D-M mutant, and 14 A-D-M WT). Unsupervised hierarchical clustering of the top 3000 variable genes across the PanNETs revealed two distinct clusters where almost all A-D-M mutant PanNETs were found in one cluster (Figure 2a). The grouping of A-D-M mutant PanNETs into one distinct cluster by gene expression was robust to the number of most variable genes used for clustering (Supplementary Fig 1). Principal component analysis (PCA) separated the A-D-M mutant PanNETs from the A-D-M WT PanNETs along the first principal component (corresponding to the component comprising the largest variation in gene expression) (Figure 2b). The separation of A-D-M mutant PanNETs from A-D-M WT PanNETs by PCA was robust to the number of top variable genes used (Supplementary Fig 2). These data show that A-D-M mutant tumors have a distinct gene expression pattern from that of A-D-M WT PanNETs. Neither hierarchical clustering nor PCA from gene expression revealed further subgrouping of the tumors with single mutations in *ATRX*, *DAXX*, or *MEN1* or double mutations in *ATRX*/*MEN1* or *DAXX*/*MEN1*.

**Figure 2.**
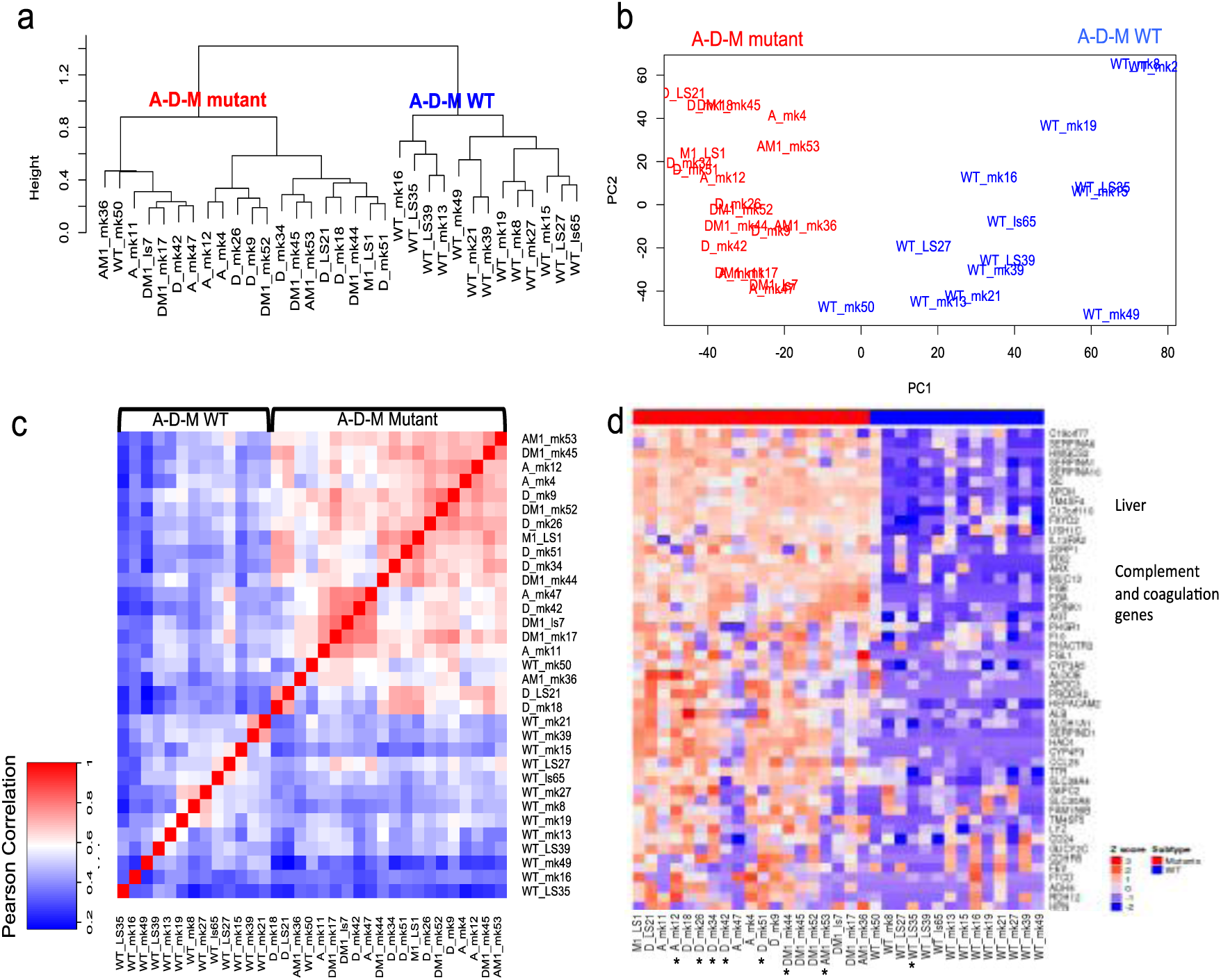
A-D-M mutant and WT PanNETs as two distinct gene expression groups. a) Unsupervised clustering of PanNETs using top 3000 variant genes across all samples revealed two distinct robust clusters. The two clusters almost perfectly separate A-D-M WT panNETs from A-D-M mutant panNETs. b) Principal component analysis using top 3000 variant genes separated the A-D-M mutant from A-D-M WT PanNETs along the first principal component (PC1). A-D-M mutant panNETs were more homogeneous in gene expression than A-D-M WT as shown by smaller variation along PC1. c) Heatmap of pair-wise Pearson correlation of panNETs using top 3000 variant genes across all samples revealed a higher correlation among A-D-M mutants as compared to A-D-M WT panNETs. Red color represents higher correlation and blue represents lower correlation. d) Heatmap of top variants genes showing liver, complement and coagulation genes highly expressed in A-D-M mutant panNETs. Star (*) below sample names represent liver metastatic samples (except for A_mk12 which is a lymph node).

In hierarchical clustering, the A-D-M mutant PanNETs formed a tighter cluster than the A-D-M WT PanNETs. In PCA, the A-D-M mutant PanNETs had smaller variance along PC1 than A-D-M WT PanNETs. Pair-wise correlation of gene expression between all PanNETs, showed a higher correlation among A-D-M mutant PanNETs as compared to A-D-M WT PanNETs (Figure 2c). Among A-D-M mutant PanNETs, mutants with the same genotype (mutations in *ATRX*/*DAXX*/*MEN1*) did not show greater gene expression correlation. These data suggest that A-D-M mutant PanNETs are a more homogeneous group compared to A-D-M WT PanNETs.

Within A-D-M mutant or A-D-M WT PanNETs groups, unsupervised clustering and PCA did not reveal differences between primary and metastatic tumors. Top 100 genes with highest variance across all samples separates mutant from A-D-M WT PanNETs and showed relatively high expression of “liver-specific” genes (*APOH, ALDH1A1, FGB, APOC3* etc.) as well as complement and coagulation pathway genes (*SERPINA1, FGA, F10, CP, MT3* etc.) in A-D-M mutant PanNETs (Figure 2d;Supplmentary Fig 3). The pathological estimate of tumor purity was over 80% for all samples of PanNETs, suggesting that the liver gene signature was not caused by liver contamination. Seven A-D-M mutant and one A-D-M WT PanNETs were from the tissue of liver metastases and they had gene expression profile most similar to the genotype group of their primary PanNET counterpart (Figure 2d). We validated the distinct gene expression signature of A-D-M mutant in a larger tumor set (47 PanNETs including the 33 PanNETs where RNA sequencing was performed) using gene expression microarray technology. The 14 additional samples are comprised of 3 A-D-M WT PanNETs and 11 A-D-M mutant PanNETs (7 *MEN1* mutant, 2 *DAXX* mutant, and 2 *DAXX/MEN1* mutant)(Supplementary Fig 4).

**Figure 3.**
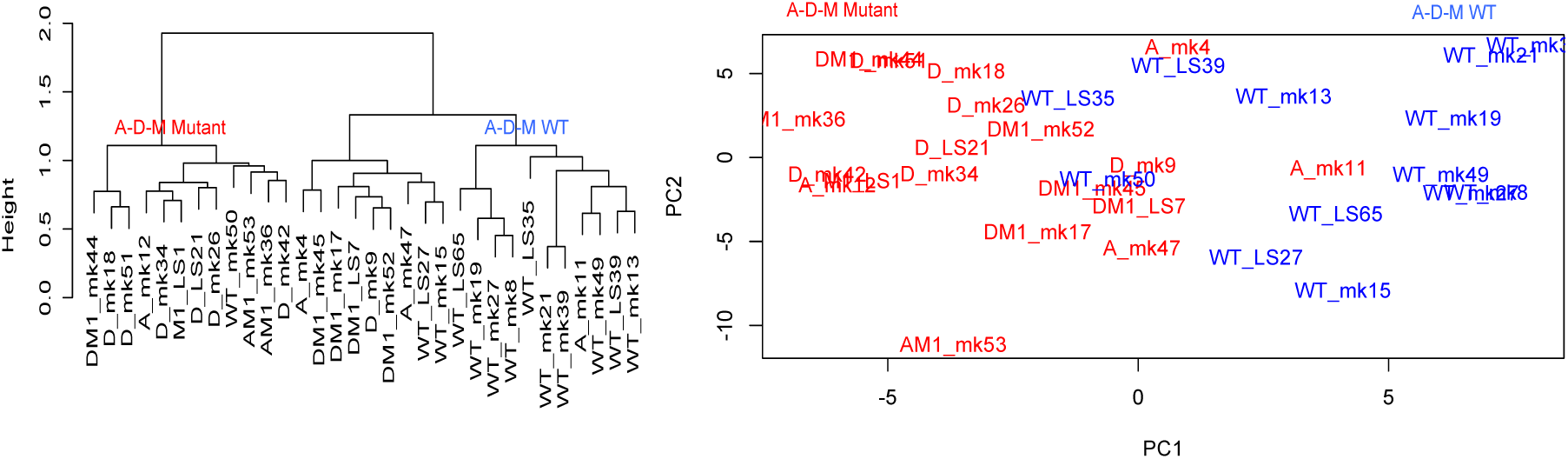
Distinct DNA methylation pattern between A-D-M mutant and A-D-M WT PanNETs. a) Unsupervised clustering of PanNETs using top 2000 variant CpG sites across all samples revealed two clusters. The two clusters separate A-D-M mutant from A-D-M WT PanNETs. b) Principal component analysis using top 2000 variant CpG sites separated A-D-M mutant from A-D-M WT PanNETs along PC1.

**Figure 4.**
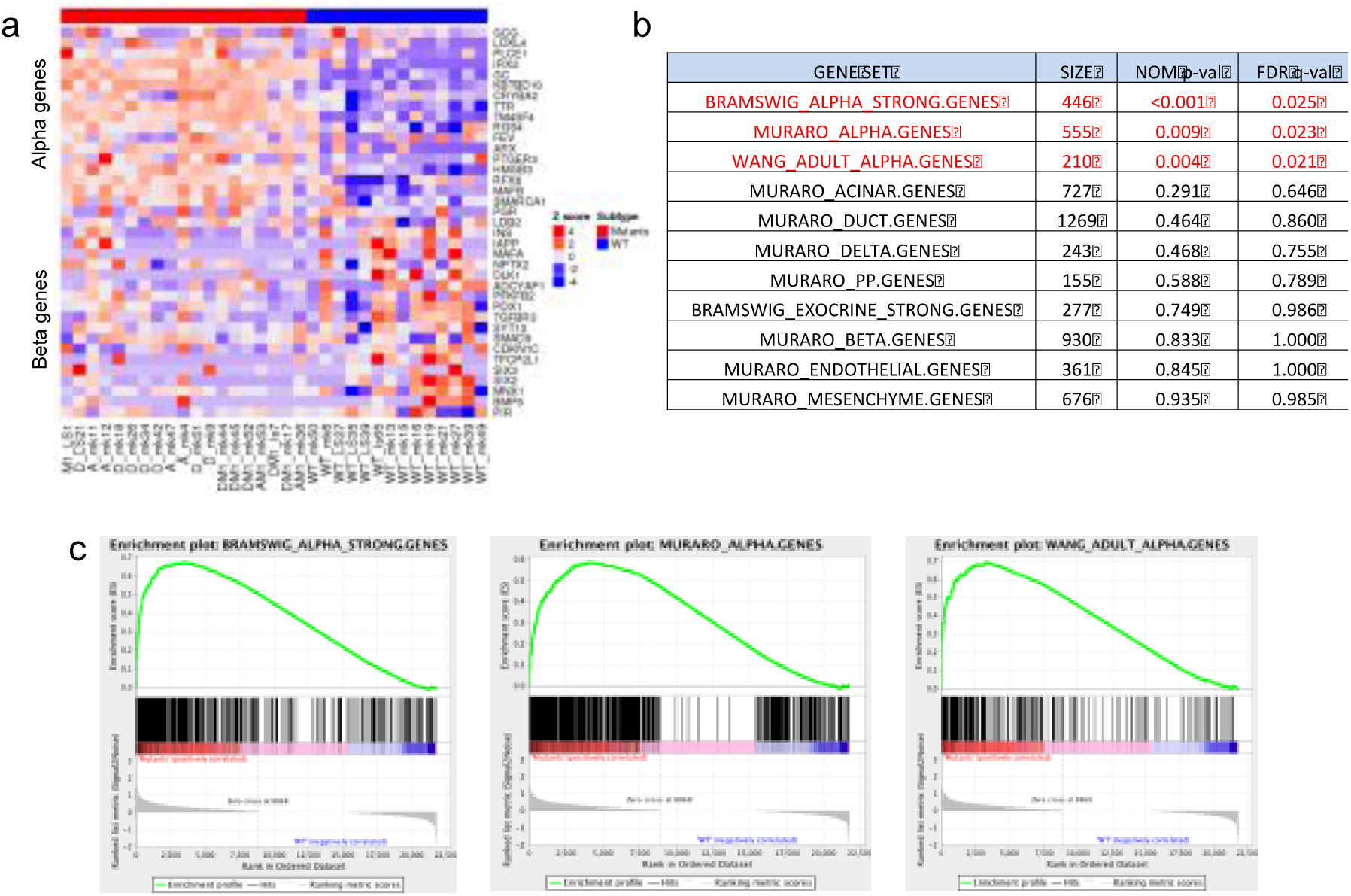
A-D-M mutant PanNETs with alpha-cell signature. a. Heatmap of gene expression for top 20 alpha and beta cell-specific genes from Muraro *et al*^6^ revealed alpha cell specific genes are highly expressed in A-D-M mutant panNETs. A-D-M WT panNETs are more heterogeneous in gene expression but some show high beta cell specific gene expression. b. Gene set enrichment analysis show A-D-M mutant panNETs to be enriched for expression of alpha cell specific genes. Pancreas cell type (alpha, beta, delta, PP, acinar, ductal) gene signatures were obtained from three different published dataset to access enrichment of cell type signatures in A-D-M mutant vs A-D-M WT PanNETs. Table represents GSEA results where size is the number of genes in gene set. All alpha cell gene sets (from three different sources) are significantly enriched in A-D-M mutant panNETs (highlighted in red). No other cell types were enriched in A-D-M mutant or A- D-M WT panNETs. c. GSEA plots of significant alpha cell signatures from Bramswig et al^9^, Wang et al^17^ and Muraro et al^6^.

To investigate epigenetic differences between PanNETs, we used the Illumina 450K chip to assay the DNA methylation at 411,549 CpG sites in 32 PanNETs. Unsupervised hierarchical clustering of the top 2000 variable DNA methylation sites across the PanNETs revealed two distinct clusters where almost all A-D-M mutant PanNETs were found in one cluster (Figure 3a). Principal component analysis (PCA) separated the A-D-M mutant PanNETs from the A-D-M WT PanNETs along the first principal component (corresponding to the component comprising the largest variation in DNA methylation) (Figure 3b). The separation of A-D-M mutant PanNETs from A-D-M WT PanNETs by PCA was robust to the number of top variable DNA methylation sites used (supplementary Fig 5). These data reveal that A-D-M mutants PanNETs have a distinct DNA methylation pattern from that of A-D-M WT PanNETs. Neither hierarchical clustering nor PCA revealed differences in DNA methylation sites between the different combinations of genes mutated among the A-D-M mutant PanNETs. Within A-D-M mutant or A-D-M WT PanNETs groups, unsupervised clustering and PCA of DNA methylation did not reveal differences between primary and metastatic tumors.

**Figure 5.**
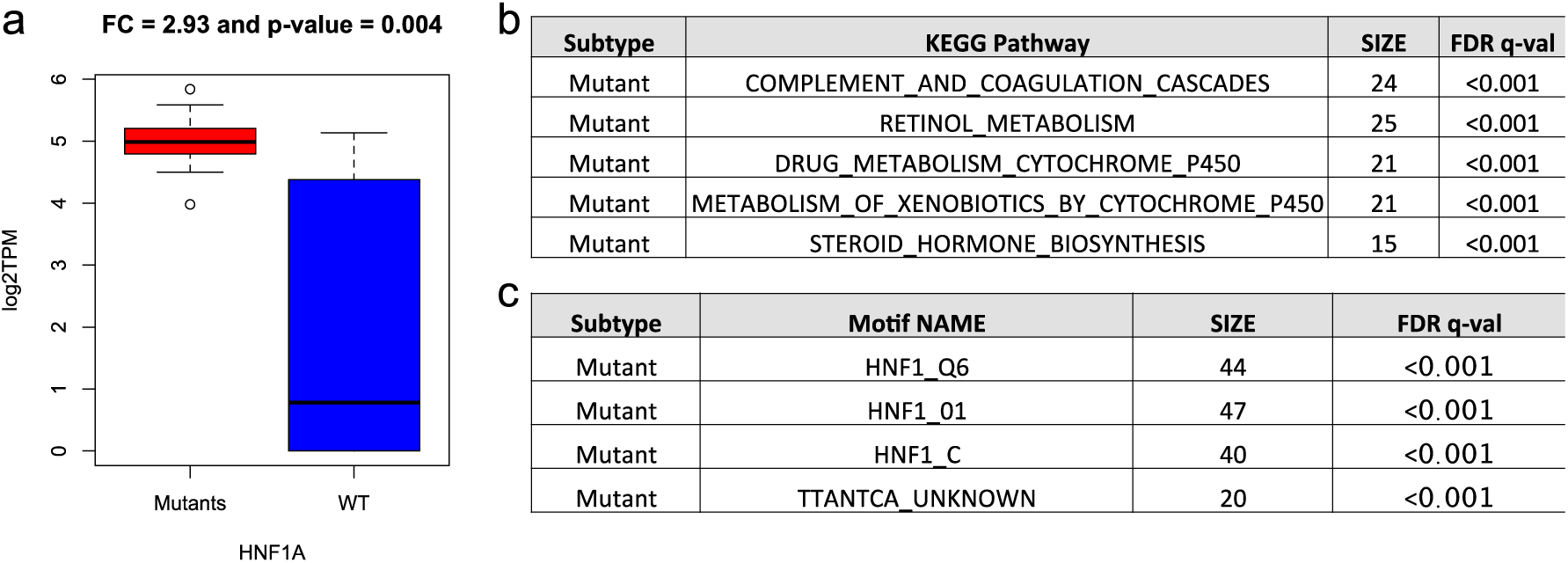
HNF1A pathway with transcriptionally up-regulation in A-D-M mutant panNETs and alpha cells. GSEA was used to find pathway enrichment from genes differentially expressed between A-D-M mutant and A-D-M WT PanNETs. a. Boxplot of HNF1A gene expression for A-D-M mutant and A-D-M WT PanNETs. HNF1A was homogeneously expressed 2.93 fold higher in A-D-M mutants PanNETs (corrected pval <0.004). b. Table represents significant KEGG pathways where genes were differentially expressed between A-D-M mutants and A-D-M WT panNETs. c. Table represents transcription factor motifs significantly enriched in promoters of genes differentially expressed in A-D-M mutants and A-D-M WT panNETs. Three HNF1 related motif gene sets from GSEA showed significant enrichment in genes over- expressed in A-D-M mutant panNETs.

### A-D-M mutant PanNET gene expression resembles that of alpha cells

There are multiple neuroendocrine cell types in the pancreas including alpha, beta, gamma, delta, and epsilon. We used gene expression data for these various pancreatic neuroendocrine and exocrine cell types from a single cell sequencing study^6^ (Supplementary Table 1) to identify gene-set signatures representing highly expressed cell-type-specific genes (Supplementary file 2). The A-D-M mutant PanNETs uniformly exhibited a gene expression signature that was very similar to that of alpha cells (Figure 4a). The A-D-M WT PanNETs were more heterogeneous in their expression of the genes among the gene set signatures for the different pancreatic neuroendocrine cell types. Greater heterogeneity of gene expression signature in A-D-M WT PanNETs was consistent with the greater heterogeneity found in global gene expression.

To further investigate the gene expression signature of A-D-M mutant PanNETs we performed gene set enrichment analysis^7^ (GSEA) on the thirteen manually curated gene sets for pancreatic endocrine and exocrine cells from a previous study. This study assessed gene expression of individual pancreatic cell types (alpha, beta, delta, PP, acinar, ductal, mesenchyme and endothelial) enriched by flow cytometry and single cell RNAseq (Supplementary Table 1). Our analysis indicates that that only the alpha cell gene was significantly enriched in A-D-M mutant PanNETs (p-value < 0.009) (Figure 4b and c)(Supplementary table 2).

Alpha and beta cell lineage specific genes were examined for the A-D-M mutant and WT PanNETs. *ARX*, *IRX2*, and *TM4SF4* were all highly expressed in A-D-M mutant PanNETs compared to A-D-M WT PanNETs (Supplementary Fig 6). Surprisingly, *GCG* (glucagon) expression was lower in A-D-M mutants as compared A-D-M WT PanNETs. For beta cell specific genes, *PDX1*, *MAFA*, *INS*, and *DLK1*, all had lower expression in A-D-M mutant PanNETs than A-D-M WT PanNETs (Supplementary Fig 6). However, these genes had much greater expression heterogeneity in A-D-M WT PanNETs suggesting that some A-D-M WT PanNETs resemble beta cells and others did not (Supplementary Fig 6).

**Figure 6:**
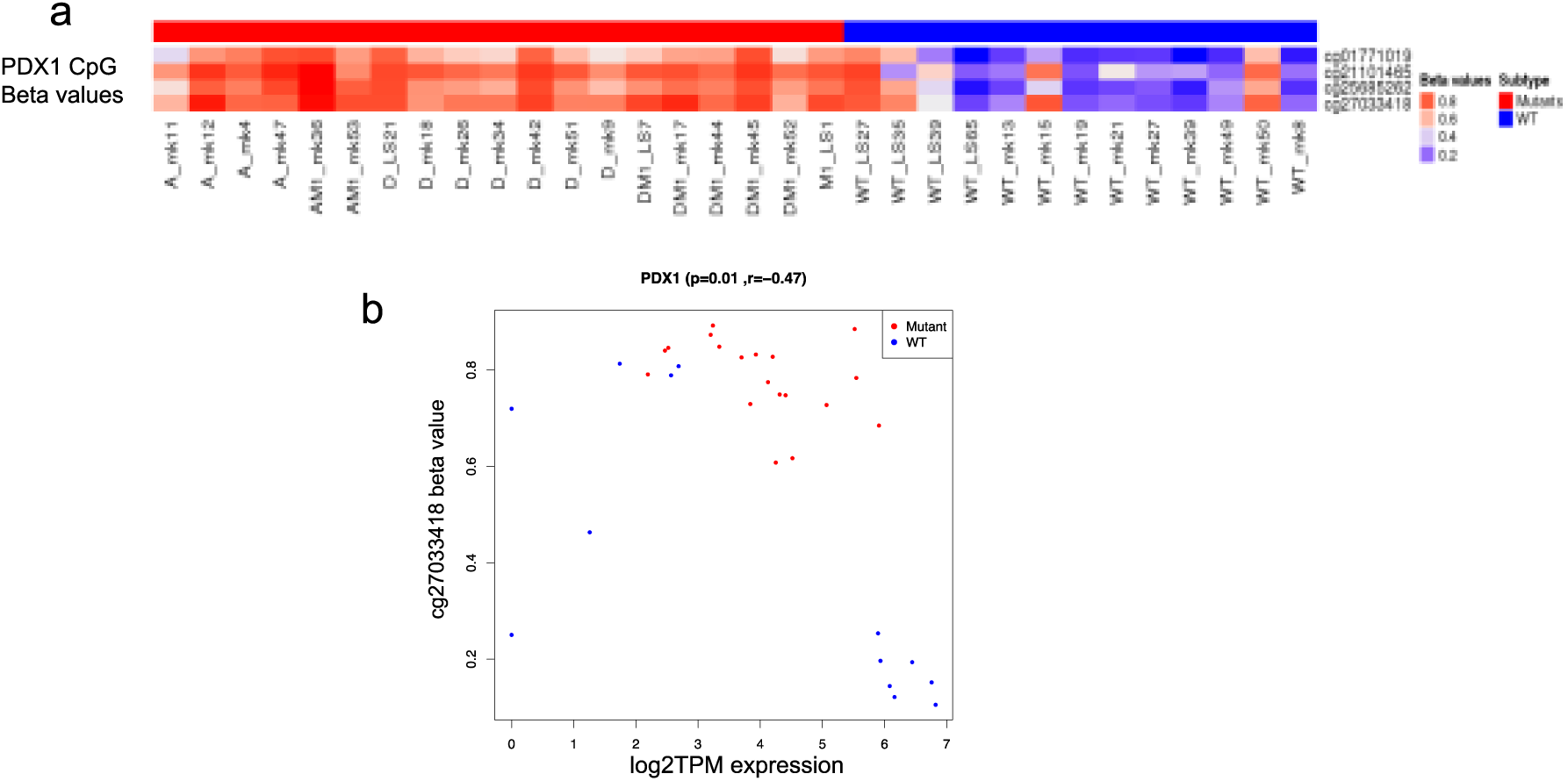
PDX1 has promoter hypermethylated and lower gene expression in A-D-M mutant panNETs. a) Four PDX1 promoter CpG sites show strong hypermethylation in A-D-M mutant PanNETs (corrected pval< 0.05). The range of beta values is from 0 to 1 and represented as blue (hypo- methylation) to red (hyper-methylation). b) PDX1 expression and promoter methylation (TSS1500 cg27033418 CpG site) across all samples showing separation of A-D-M mutant and A-D-M WT PanNETs.

### HNF1A pathway is transcriptionally up-regulated in A-D-M mutant PanNETs and alpha cells

HNF1A is one of the most significantly differentially expressed genes between A-D-M mutant and WT PanNETs. HNF1A is a homeobox family transcription factor that is highly expressed in the liver and is involved in the regulation of several liver-specific genes. The expression of HNF1A was 2.93 fold higher in A-D-M mutant PanNETs than A-D-M WT PanNETs (corrected p-value < 0.004) (Figure 5a). Differentially expressed genes (DEgenes) between the A-D-M mutant and A-D-M WT PanNETs were found in 1478 genes (with greater than 3 fold change and corrected p-value < 0.05, See Methods section)(Supplementary file 3). Functional pathway enrichment for DEgenes using preranked GSEA revealed the complement and coagulation cascades, retinol metabolism, and drug metabolism to be up-regulated in A-D-M mutant PanNETs (see Methods section) (Figure 5b; Supplementary file 3). The differentially expressed genes were also enriched for HNF1A transcription factor motifs in their promoters (FDR < 0.001, Figure 5c). The complete list of significant TFs motif is present in Supplementary file 3. Taken together, the A-D-M mutant PanNETs had higher expression of *HNF1A* along with many of its transcriptional target genes associated with liver function. In addition, the transcriptional regulator of *HNF1A, HNF4A*^*8*^ was expressed 3.02 fold higher in A-D-M mutant PanNETs (p-value < 0.009). We used gene expression data from Bramswig et al^9^ to show *HNF1A* expression was increased in alpha cells compared to beta cells (p-value < 0.008), and the 465 alpha cell specific genes in the pancreas were enriched for transcriptional targets of *HNF1A* and for having *HNF1A* TF motif in their promoters (See Methods section; Supplementary Table 3).

Many of the most differentially expressed genes and highly expressed in A-D-M mutant PanNETs are targets of *HNF1A* and are involved in protein secretion, transport and metabolism (*APOH, ALB, AFM, HAO1, UGT1A3, UGT1A1, GC, G6PC, TM4SF4, PKLR* etc). APOH is expressed 8.46 fold higher in A-D-M mutant PanNETs (p-value < 10^-5^) and is potentially a good diagnostic biomarker for A-D-M mutant PanNETs. We perform IHC staining for APOH and show positive staining in 70±2.5% of A-D-M mutant and only 18±2.0% of A-D-M WT PanNETs (Supplementary Fig 7 and Supplementary file 1).

### The *PDX1* gene is more highly methylated and expressed at lower levels in A-D-M mutant PanNETs

DNA methylation differences between the A-D-M mutant and A-D-M WT PanNETs were found at 378 CpG sites (corrected p-value < 0.05 and difference in beta value > 0.2, see Methods section), 287 of which were found in genes and 91 in intergenic regions (Supplementary file 4). Of the 287 differentially methylated genic CpG sites, 70 (associated with 59 genes) were found between 1500 bp 5’ to transcriptional start site (TSS) and 200 bp 3’ to TSS or within first exon, a region where DNA methylation is associated with transcriptional repression^10^. Thirteen of the 59 genes were also found to be differentially expressed (with fold change greater than 3 and corrected p-value < 0.05, see Methods section) and seven genes that were hypomethylated in A-D-M mutant and over-expressed are *APOH, CCL15, EMID2, PDZK1, HAO1, BAIAP2L2, and NPC1L1.* One gene*, TACR3*, was hypomethylated in A-D-M WT and over-expressed (Supplementary file 4). Four of the 70 CpG sites were found in the gene *PDX1* (pancreatic and duodenal homeobox 1), a transcription factor necessary for pancreatic development and beta cell maturation. *PDX1* functions in the cell fating of endocrine cells, favoring the production of insulin positive beta cells and somatostatin positive delta cells while repressing glucagon positive alpha cells^11^. These four CpG sites were all hypermethylated in A-D-M mutant PanNETs (Figure 6a) and the expression of *PDX1* was 2.92 fold higher in A-D-M WT PanNETs (p-value < 0.005) (Figure 6 a and b). In contrast, while *ARX* was highly expressed in A-D-M mutant PanNETs compared to A-D-M WT PanNETs, the promoter and first exon of ARX are not differentially methylated.

## Discussion

Here, we found that A-D-M mutant PanNETs form a distinct subgroup on the basis of their gene expression profile and DNA methylation pattern. Moreover, this subgroup is more homogeneous based on gene expression profile than the A-D-M WT PanNETs. The gene signature of the A-D-M mutant PanNETs strongly corresponds to the genes that are specifically expressed in alpha cells including genes known to define alpha cells such as ARX and “liver-specific” genes such as HNF1A and its transcriptional targets. Conversely, PDX1, a gene critical to the beta cell lineage is transcriptionally repressed in A-D-M mutant PanNETs and the PDX1 promoter is hypermethylated. On the other hand, WT PanNETs have heterogeneous gene expression profiles and their gene mutational landscape is less understood.

The pancreas is comprised of many different cell types including acinar, ductal, and at least five neuroendocrine cell types including alpha, beta, gamma, delta, and epsilon cells. There are two plausible explanations for the “alpha cell-like” expression pattern of A-D-M mutant PanNETs. Either an alpha cell or an uncharacterized cell type with an alpha-cell like gene expression profile is the cell-of-origin for PanNETs with mutations in ATRX, DAXX and MEN1. Alternatively, loss of ATRX, DAXX, or MEN1 genes may promote pancreatic neuroendocrine (or progenitor) cells types to reprogram their gene expression profiles to resemble alpha cells. Although, it remains unclear whether there are pancreatic stem or progenitor cells in adult pancreas.

ATRX-DAXX and MEN1 are involved in distinct biochemical pathways to regulate gene expression. Therefore, we would expect that loss of these proteins during transformation of A-D-M mutant PanNETs would result in a more heterogeneous gene expression profile. Due to the high degree of homogeneity of the A-D-M mutant PanNETs at the level of gene expression and the strong expression of genes that are known to be alpha cell specific, we hypothesize that alpha cells are the cell-of-origin for this group of tumors. In addition, *MEN1* and *ATRX*/*DAXX* mutations occur alone or in a combined pattern suggest that they have independent oncogenic activities in A-D-M mutant PanNETs, making the idea of reprogramming to a homogeneous alpha-like cell state less probable. Some of the A-D-M WT PanNETs have a strong beta cell signature and these may have arisen from beta cells (Figure 4a). However, other A-D-M WT PanNETs have neither alpha nor beta cell signatures, which may arise from other cell types in the pancreas.

Conditional knockouts of MEN1 in mice support the model of an alpha cell origin for A-D-M PanNETs^13^,^14^. The restricted deletion of MEN1 to alpha cells surprisingly led to the development of insulinomas^14^. Most of our A-D-M mutant PanNETs were nonfunctional (26 of 33 PanNETs) but the functional tumors were insulinoma and VIPoma, even though their gene expressions have alpha cell signature. In other mouse models of PanNETs^12^,^13^, MEN1 deletion using the insulin or PDX1 promoter driven Cre construct, insulinomas and well differentiated PanNETs were also observed. However, Cre expression may be leaky in these models and further study is needed to understand the heterogeneity of the cells in the tumors that develop and trace the cell of origin or transdifferntiated state of the cancer cells.

In our gene expression analysis, we have not identified the oncogenic pathways activated in A-D-M mutant PanNETs. MEN1 has been shown to upregulate expression of long noncoding RNA MEG3 in MIN6 mouse insulinoma cell line^15^. In the same study, they show MEG3 represses expression of the oncogene MET leading to delayed cell cycle progression and reduced cell proliferation. In a different study, MEN1 and DAXX were shown to repress the expression of the membrane metalloendopeptidase (MME) and mutations in MEN1 or DAXX result in loss of this repression leading to neuroendocrine tumor proliferation^16^. Our data is consistent with these studies when comparing A-D-M mutant to WT PanNETs, showing that A-D-M mutant PanNETs have lower expression of MEG3 (7.3 fold lower, p-value < 4.3E-07), higher expression of MET (3 fold higher, p-value < 0.003), and higher expression of MME (4 fold higher, p-value < 0.001). Among A-D-M mutant PanNETs, we do not see expression differences of MEG3, MET, and MME depending on mutation status of ATRX, DAXX, and MEN1.

While PanNETs may seemingly represent as a single clinical disease, they can be further characterized into different subtypes based upon their cell lineage and the associated molecular genotype. Understanding these biologic and molecular mechanisms of PanNETs may explain the unpredictable outcome of the disease and facilitate the development of unique and targeted therapeutic strategies.

## Method and Materials

### Patient’s information

Retrospective and prospective reviews of well-differentiated, pancreatic neuroendocrine neoplasms were performed using the pathology files and pancreatic database at MSKCC with IRB approval. All patients were evaluated clinically at our institution with confirmed pathologic diagnoses, appropriate radiological and laboratory studies, and surgical or oncological management. Follow-up information was obtained for all cases.

### Tissue acquisition and nucleotide extraction

Briefly, cases of pancreatic neuroendocrine tumors were identified. Fresh-frozen tumor and paired normal tissues were obtained from MSKCC’s tissue bank under an Institutional Review Board protocol. Histopathology of all tissues was evaluated on hematoxylin and eosin stained sections by an experienced gastrointestinal-hepato-pancreatobiliary pathologist to insure the nature of the tissue, greater than 80% tumor cellularity and absence of necrosis. The relevant tissues were then macro-dissected (20-25 mg) and DNA/RNA extraction using Qiagen’s DNeasy Blood & Tissue Kit and RNeasy Mini Kit, respectively was carried out according to the manufacturer’s protocols (Qiagen, Valencia, CA).

### Sanger sequencing for gene mutation

All exons of the DAXX, ATRX, and MEN1 genes were amplified by PCR and then sequenced using Sanger sequencing. Every mutation detected was validated by bidirectional Sanger sequencing on the tumor-normal pairs. To maintain the correct sample annotation, we used mutation status as sample name with sample ID (For example, A_mk11 sample is ATRX mutant and mk11 is sample ID). Supplement file 1 contains all the clinical information, mutational profile and sample annotation. Online Oncoprint were used to plot create figure1a.

### PanNETs Transcriptome Sequencing and data analysis

RNA Library preparation and RNA sequencing was done by MSKCC Genomics Core Laboratory using Illumina HiSeq with (2 x 75 bp paired end reads) to a minimum depth of ~ 50 million reads were generated for each sample. Raw fastq files were probed for sequencing quality control using FastQC [http://www.bioinformatics.babraham.ac.uk/projects/fastqc]. Sequencing reads were mapped to human transcripts corresponding to Genepattern^18^ genome (hg19 version) GTF annotations using RSEM with default parameters. RSEM package^19^ were used to prepare the reference genome with given GTF and calculated expression from mapped BAM files. STAR^20^ aligner was used to map reads in RSEM algorithm. Transcripts mapped data were normalized to TPM (Transcript Per Million) from RSEM and log2 transformed. This log2TPM values were used for all downstream analysis. Unsupervised clustering and Principal Component analysis was conducted to elucidate subtypes structure using top 3000 variant genes as input. To query robustness of this subtyping, multiple variant gene sets were used and repeated the same process of unsupervised clustering. Top 100 variable genes were used to find genes, which were highly expressed in each subtype. Subset of these genes is selected to show in figure 2d for liver and complement system genes. To find differentially expressed genes (DEgenes) between mutant and wild-type panNETs, we used DeSeq2 R package^21^ on raw count (values from RSEM). We used significance cutoff with greater than 3 fold change and corrected p-value < 0.05 to call a gene as DEgene. GSEA Preranked^7^ method was used on DEgenes to find significant KEGG pathways, motif and biological process.

### Clustering and Principal Component Analysis

For unsupervised clustering on log2TPM, we used Pearson distance metric and ward.D2 hclust method(unless stated otherwise). PCA analysis was done using *prcomp* in R. R (http://www.r-project.org/) was used for all the analysis and visualization of data.

### Pancreatic Endocrine and Exocrine gene set (PEEGset) from published dataset^6,9,17^

The neuroendocrine cells in the pancreas include alpha, beta, delta, pancreatic polypeptide (pp)- producing and vasoactive intestinal peptide (VIP)-producing cells. Gene sets representing different endocrine islet and exocrine pancreatic cells (PEEGset) were obtained from three metadata^6^,^9^,^17^ (supplementary Table 1). We created 13 PEEGset representing all major cells from endocrine and exocrine pancreases. Supplementary Table 1 shows these gene sets with major cell types and number of genes in each set. These gene set were used as prior defined gene set for GSEA analysis.

## Gene Set Enrichment Analysis on Major Islets cell types

Gene Set Enrichment Analysis^7^ (GSEA) was performed on the log2TPM expression values of all samples using downloaded version of GSEA software (Broad Institute, Cambridge, MA, USA) to identify the statistically enriched gene sets between A-D-M mutant and A-D-M WT PanNETs. Published pancreatic islet endocrine and exocrine cells signatures were used as prior defined sets as an input. We used all default parameters to perform GSEA on this gene sets to determine the enrichment of specific cell signature enrichment in the panNET subtypes. We ran GSEA on 1000 permutation mode on phenotypic label to generate FDR and enrichment score (ES) for each gene set. Significant gene set was filtered based on FDR q-values (cutoff of 0.05).

## Bramswig et al^9^ FACs sorted normal alpha and beta cells gene expression

We extensively used Bramswig et al^9^ FACs sorted RNAseq data to understand normal alpha and beta cells and correlated their gene signature sets with our A-D-M mutant and A-D-M WT panNETs. We downloaded supplement file for total RNA seq normalized expression data for alpha (3 replicate) and beta (3 replicate) and exocrine cells (2 replicate). Bramswig et al^9^ provide strong genes associated with alpha, beta and exocrine cells as supplement file. We used this strong cell specific genes and created gene set for alpha, beta and exocrine and named as Bramswig_et_al gene set. HNF1A gene expression values were fetched to check whether HNF1A is over expression in normal alpha as compared to beta and exocrine. We applied Student ttest’s between three alpha and three beta samples to calculate p-value for HNF1A gene expression. Bramswig et al strong alpha cell genes (n=465) were queried to check for HNF1A transcription factor motif enriched using online GSEA version (C3 TFs motif database).

## Microarray Gene expression analysis

RNA extracted from PanNETs samples were submitted for Affymetrix microarray profiling using chip hgu133a2 array. All following analysis was done in R. Briefly, the raw Affymetrix CEL files were loaded in R (simpleaffy^22^). Normalized expression values were calculated by the GC Robust Multi-array Average (GCRMA^23^) algorithm and subjected to mean transformation in order to collapse all probes to respective genes in R using collapseRow^24^. We then followed the same clustering and PCA procedure that were done on RNAseq expression data. Collapsed average gene expression values were imported in GSEA to run against 13 PEEGset cell types to find enrichment.

## 450K DNA methylation array analysis

DNA extracted from PanNETs samples and interrogated using the Illumina 450K platform (Illumina Inc. San Diego, CA) to access the DNA methylation profiles. All the analysis was performed using ChAMP^25^ version 2.6.0 open source software implemented in R. Briefly, IDAT file raw data were imported in R and filtered to exclude samples with detection p-value <0.01 and beadcount <3 in at least 5% of samples and normalized using FunctionNormalization^26^. This normalization method correct for background; remove dye bias followed by Quantile normalization. Unsupervised clustering and PCA were done on top variants 2000 probes (Var2000) across all samples to find classes of PanNETs. We repeated this clustering using different number (Var10000, Var5000, Var3000, Var1000 and Var500) of probes to check robustness of this subtyping. Differentially methylated CpG sites (DMP) between the A-D-M mutant and A-D-M WT PanNETs were identified using champ.MVP using the all default parameter method (Bonferroni-Hochberg) to adjust the p-value(<0.01). Significant DMP sites from respective genes were compared to DEgenes to find overlapping dysregulated genes in each subtype.

## Immunohistochemistry (IHC)

A representative, formalin-fixed, paraffin-embedded tissue section (4 μm thick) of each case was submitted to our institution’s core facility to perform immunohistochemistry-using antibodies recognizing the APOH proteins. Briefly, sections were de-paraffinized and pre-treated in Cell Conditioning 1 (CC1 mild; Ventana Medical Systems, AZ, USA) using an automated staining system (Ventana Discovery XT Autostainer; Ventana Medical Systems Inc, Tucson, AZ). Primary antibodies were applied for 60 min at a dilution of 1:100 for APOH (anti-APOH, polyclonal antibody; Proteintech). The sections were then incubated for 60min with secondary antibody (1:200) followed by DAB Map detection (DAB visualization; Ventana Medical Systems). Cytoplasmic (APOH) labeling in at least 50% of the tumor cells was considered positive. In the case of APOH, normal liver tissue was used as a positive control in each experiment.

## Statistical Analysis

Data are represented as mean ± standard deviation. GraphPad Prism 6 (GraphPad Software Inc, La Jolla, Ca) was used for statistical and survival analyses. Survival analysis p-values (2-sided) were based on log-rank tests. Significance was defined as P < 0.05.

## Acknowledgements

This research was supported by Starr Cancer Consortium Grant (CDA and LHT), Raymond and Beverly Sackler Foundation (LHT), Caring for Carcinoid Foundation (CDA and LHT), Mushett Family Foundation (LHT), MSKCC Support Grant/Core Grant (P30CA008748) and the Biomedical Informatics shared resource of Rutgers Cancer Institute of New Jersey (P30CA072720).

## Supplementary Files, Tables and Figures

### Supplementary Files for Chan S C and Laddha SV et al

*ATRX*, *DAXX* or *MEN1* mutant pancreatic neuroendocrine tumors are a distinct “alpha-cell signature” subgroup

### Supplementary Files

**Supplementary File 1:** PanNETs clinical information

**Supplementary File 2:** Gene list for each Pancreases Endo and Exocrine Gene Set (PEEGset)

**Supplementary File 3:** DEseq2 Differentially Expressed Genes, KEGG Pathways and Motif TFs

**Supplementary File 4:** Differentially Methylated Probes and PDX1 significant probes

**Supplementary File 5:** RNAseq log2TPM expression matrix for 33 PanNETs samples

---

**Supplementary Figure 1:**
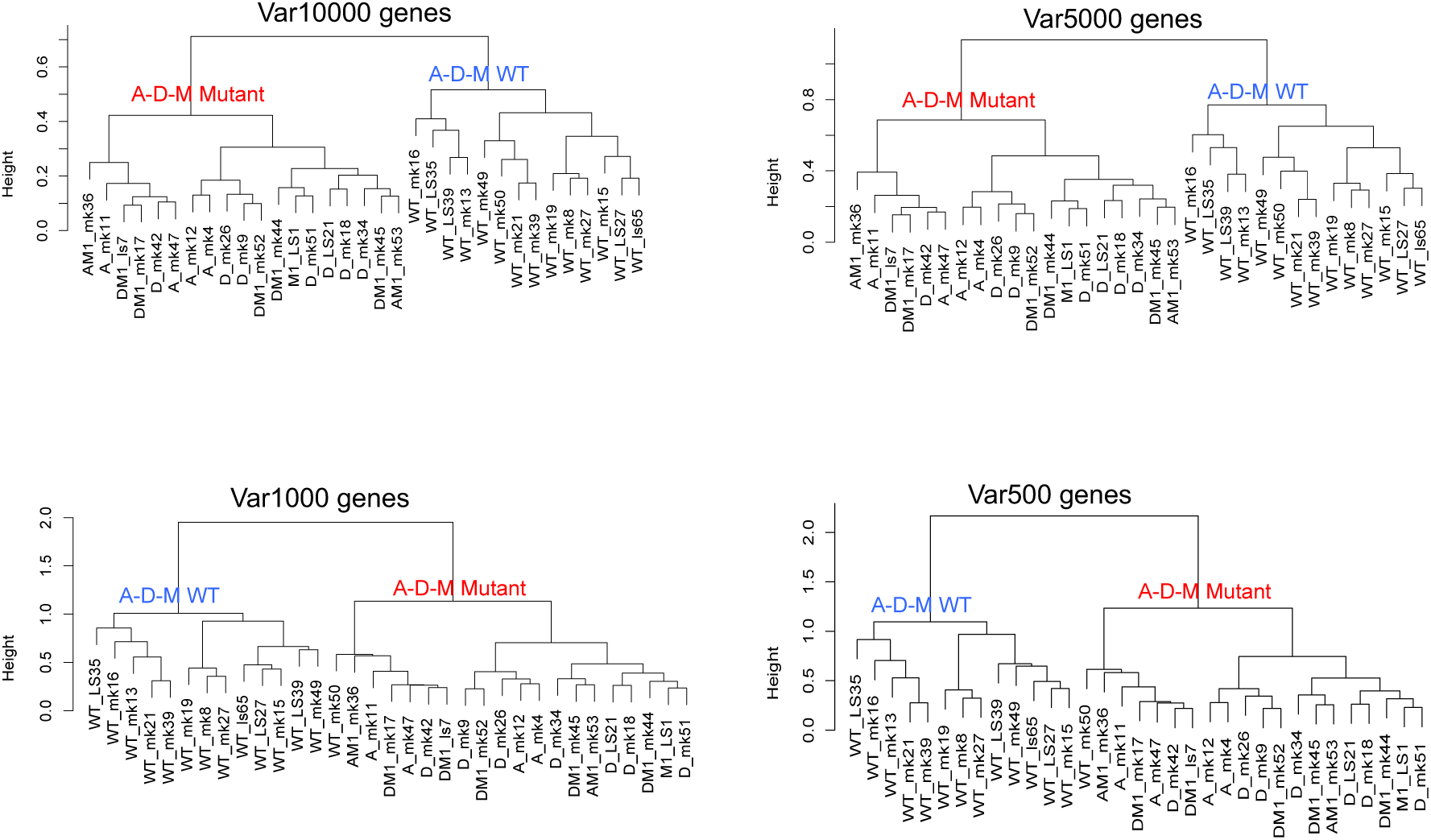
Unsupervised clustering of top variable (Var) number of genes for four different gene set (Var10000, Var5000, Var1000 and var500) shows robust subtyping. Var10000 is a gene set with 10000 top variable gene based on gene expression across all samples. All four gene set show robust stratification of A-D-M mutant (in red) from A-D-M WT (in blue) PanNETs.

**Supplementary Figure 2:**
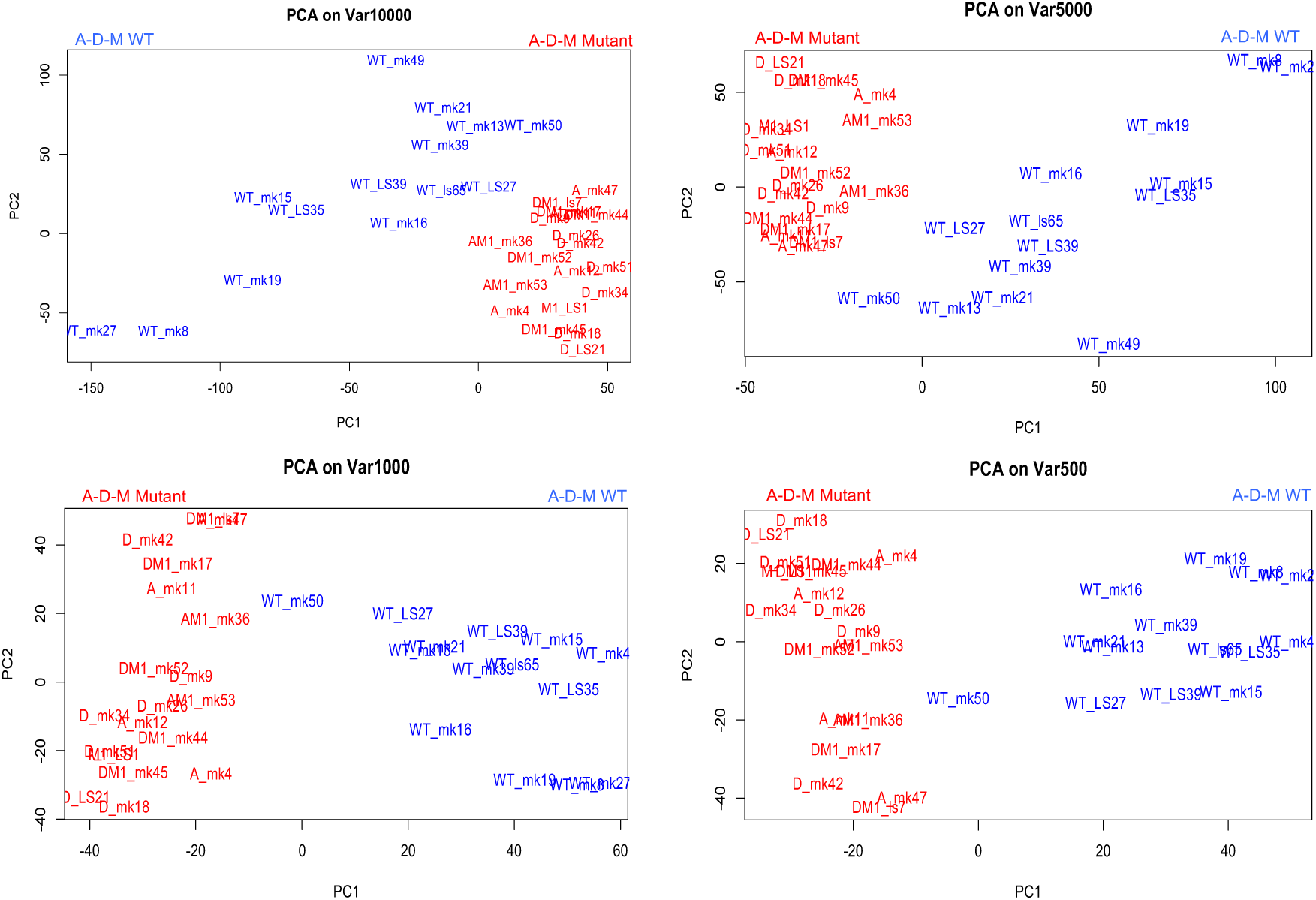
Principal Component analysis of top variable (Var) number of genes for four different gene set (Var10000, Var5000, Var1000 and var500) shows robust subtyping. A-D-M mutant are highlighted in red and A-D-M WT panNETs in blue.

**Supplementary Figure 3:**
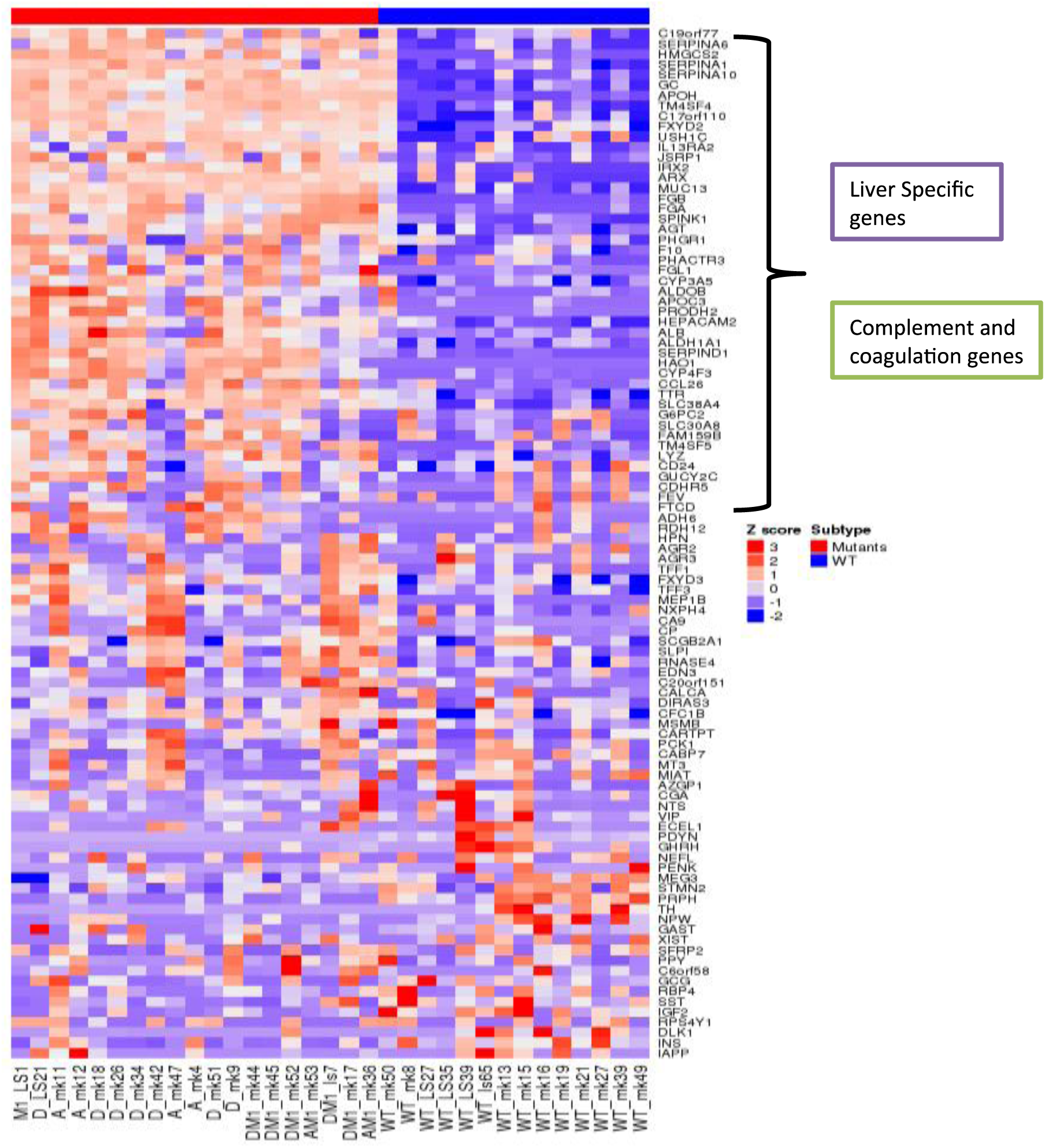
Heatmap of top 100 variance genes stratify A-D-M mutant from A-D-M WT panNETs. Many of liver and complement specific genes are highly expressed in A-D-M mutants.

**Supplementary Figure 4:**
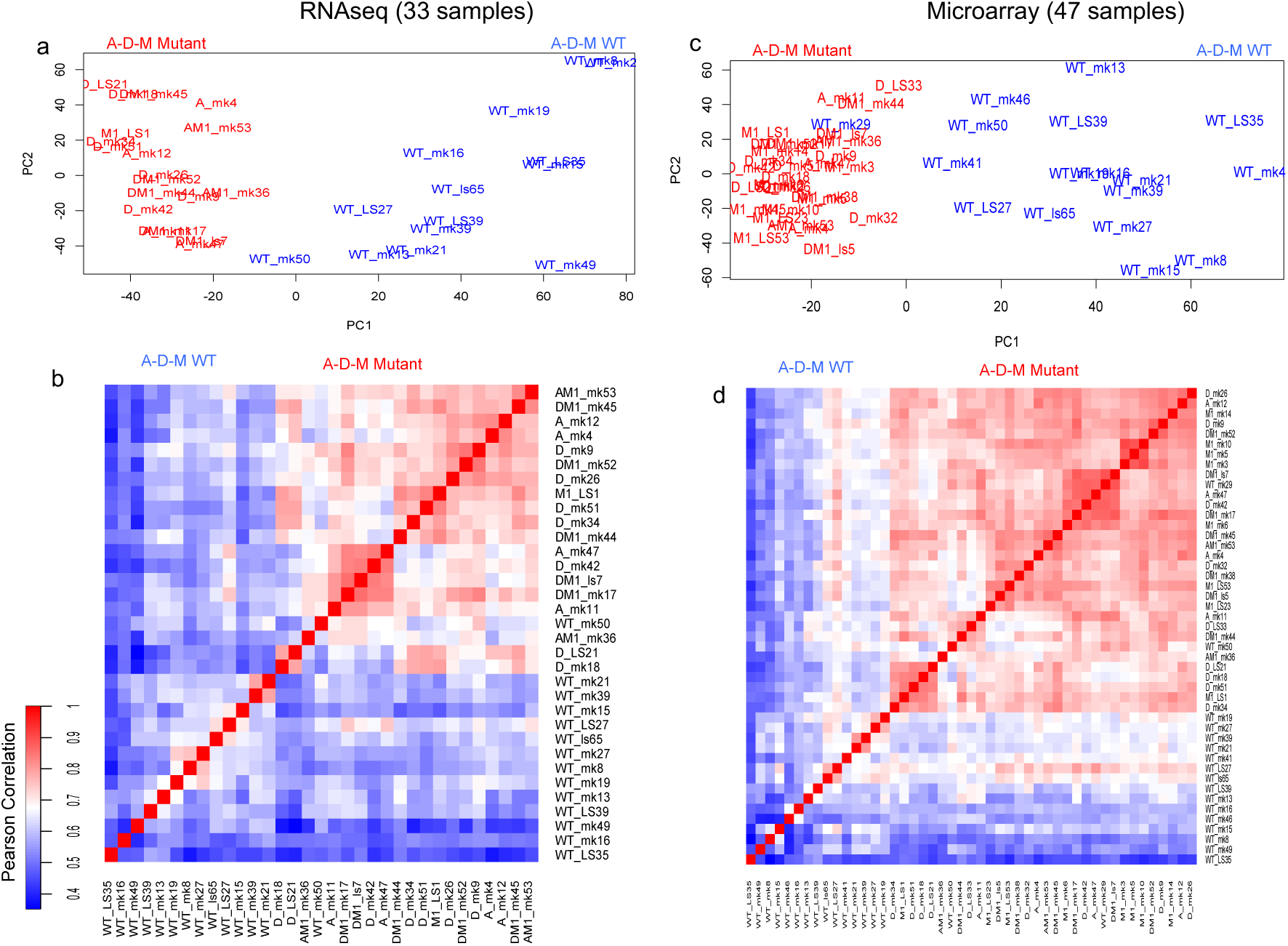
Validation of subtyping in a larger PanNETs set (n=47). The distinct gene expression signature and greater degree of homogeneity of A-D-M mutant over A-D-M WT PanNETs were validated in a larger tumor set (47 PanNETs including the 33 PanNETs where RNA sequencing was performed) using gene expression microarray technology (a and b are PCA and Pearson correlation on Var3000 genes from RNAseq and c and d are for Var3000 genes from Microarray.

**Supplementary Figure 5:**
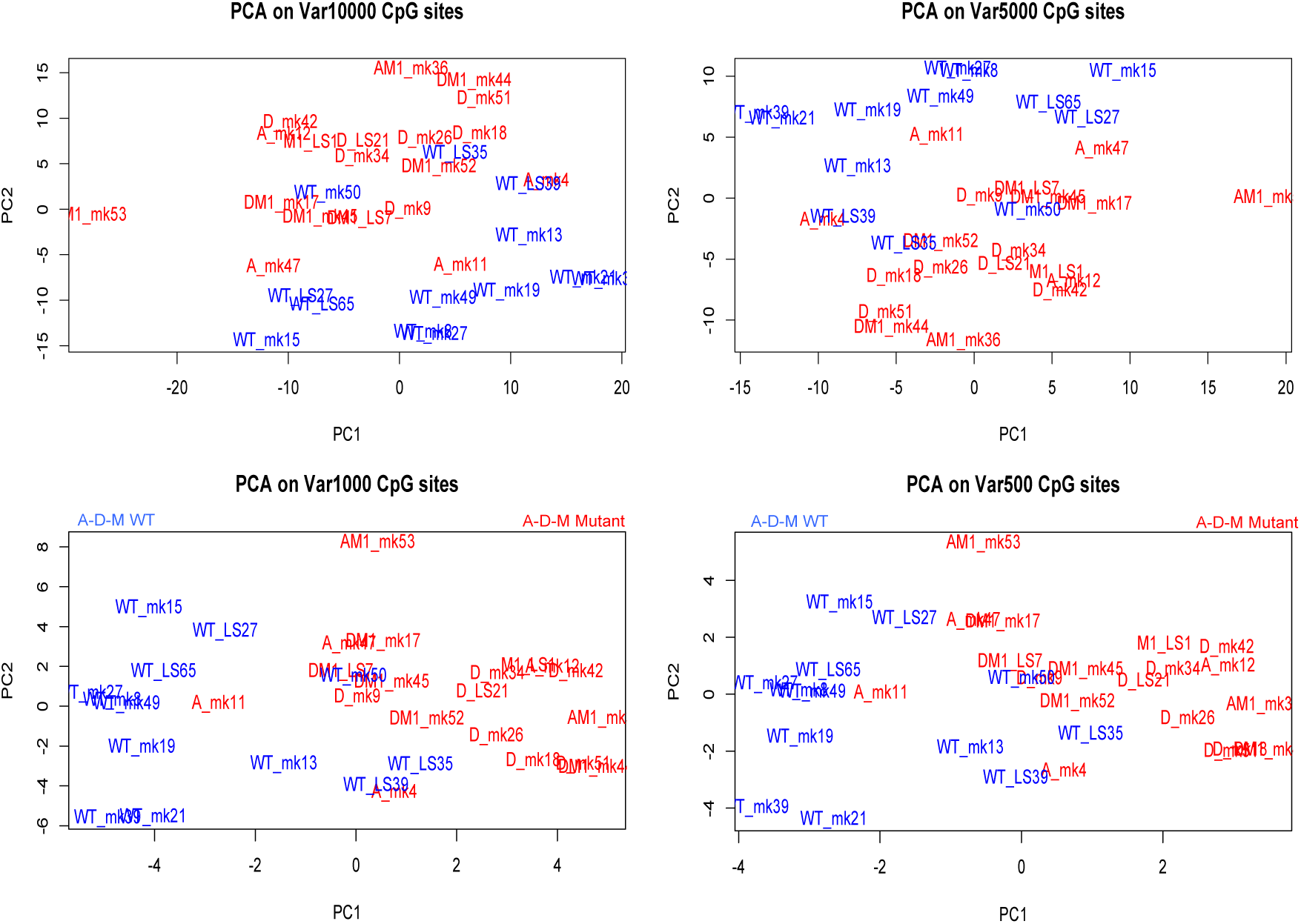
PCA of top variable number of CpG sites (450K methylation) for four different gene set (Var10000, Var5000, Var1000 and var500). A-D-M mutant are highlighted in red and A-D-M WT panNETs in blue.

**Supplementary Figure 6:**
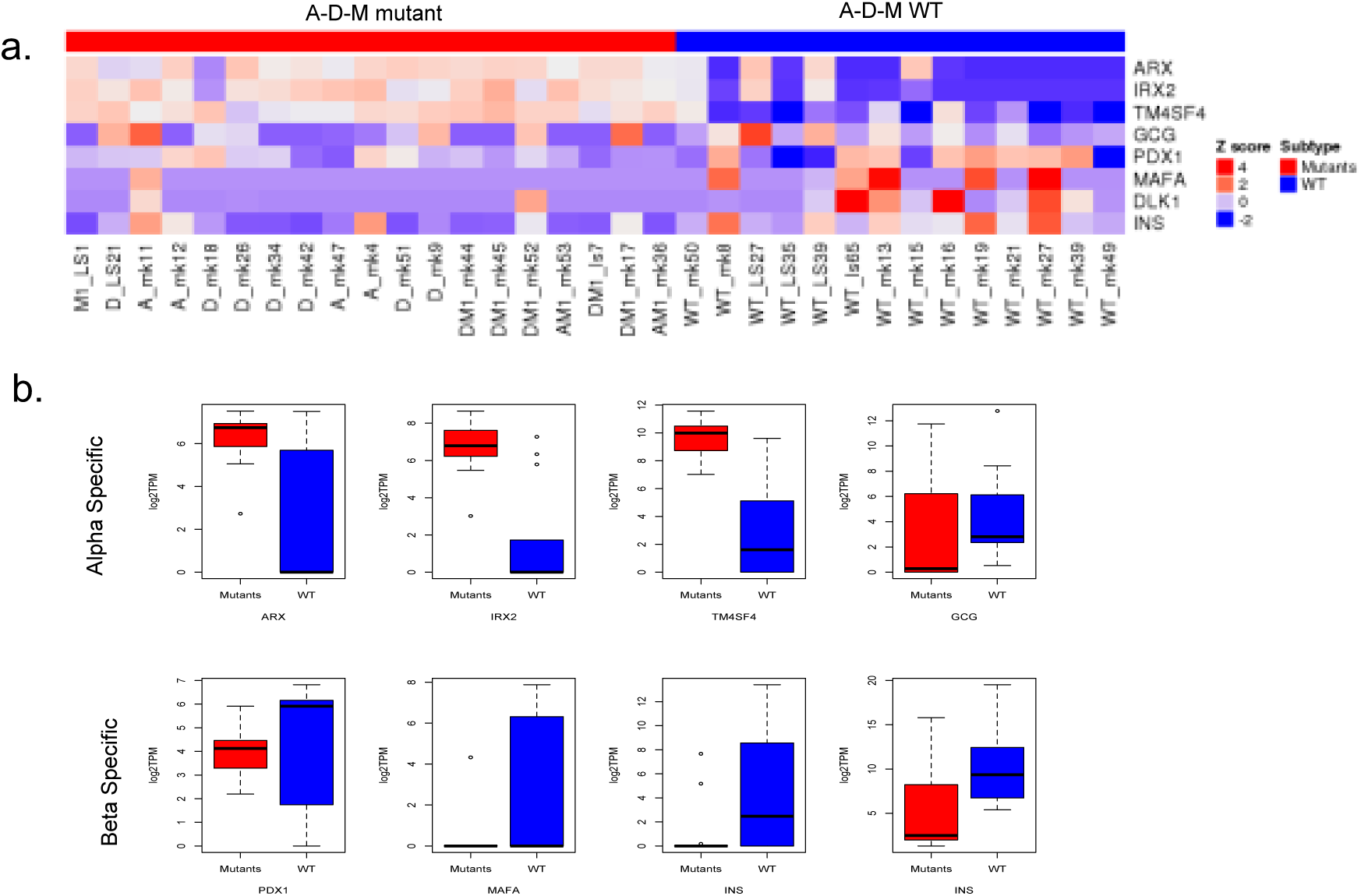
Alpha and Beta cell lineage specific genes expression and boxplot for PanNETs subtypes. Alpha Specific genes are *ARX* (TF), *IRX2* (TF), *TM4SF4* (Alpha cell surface marker), GCG and beta Specific genes are *PDX1* (TF), *MAFA* (TF), *INS*, *DLK1*. TFs represent Transcription Factor.

**Supplementary Figure 7:**
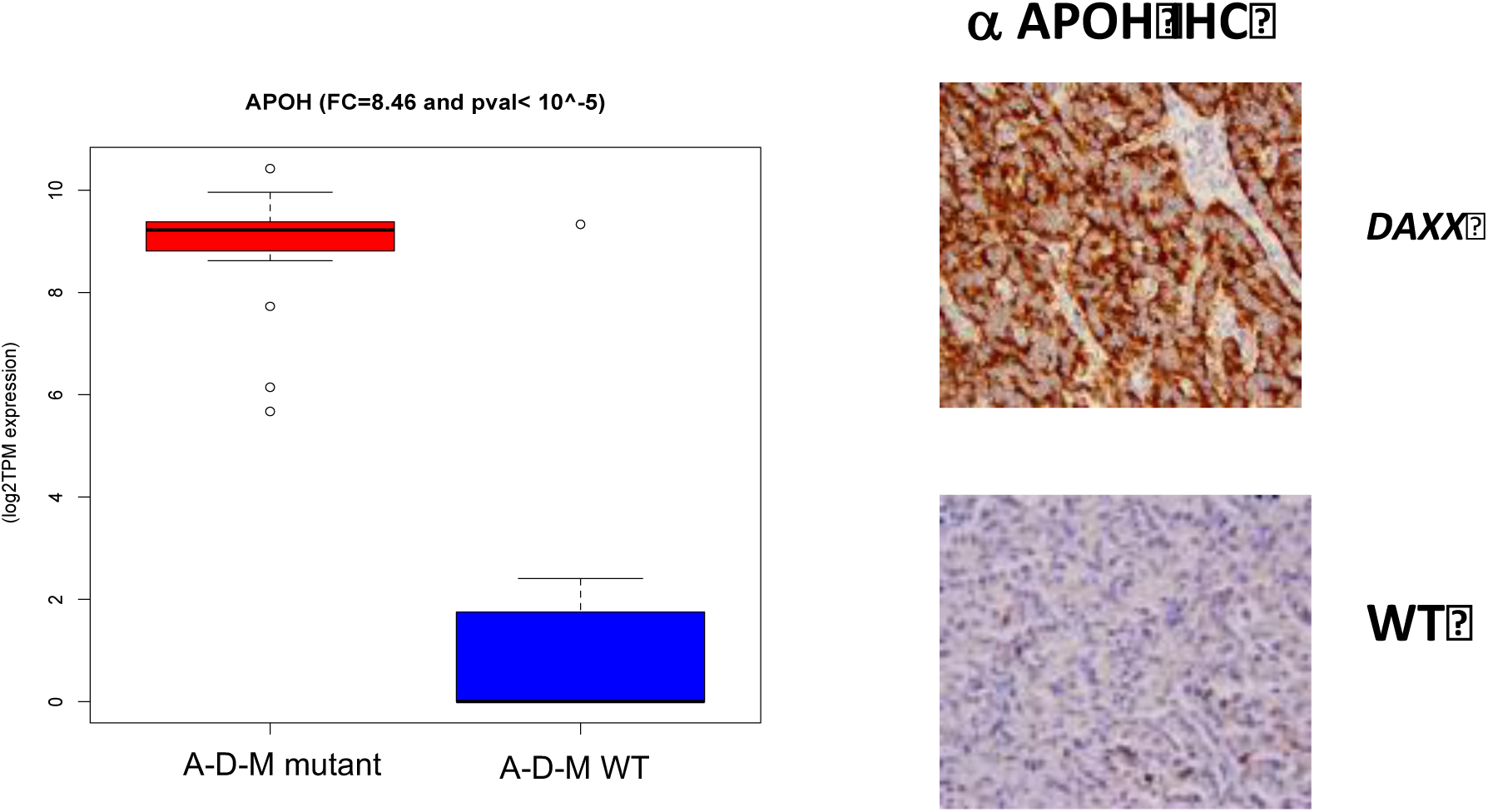
APOH gene expression and IHC. A-D-M mutant (red) has higher expression of APOH at mRNA level and Protein level. In consistent with the gene expression profile, APOH protein was strongly expressed in 70±2.5% of mutants and only18±2.0% of WT PanNETs.

**Supplementary Table1:** Pancreatic Endocrine and Exocrine Gene Set from three meta dataset. Published literature from Single cell RNA sequencing and FACs sorted total RNAseq data has been used for create PEEGset. In this study, we used following paper’s data : PMID:27364731 - Single cell transcriptomics of the human endocrine pancreas PMID:27693023 - A Single-Cell Transcriptome Atlas of the Human Pancreas PMID:23434589- Epigenomic plasticity enables human pancreatic α to β cell reprogramming

| Gene Set | PMID | Description | # of Genes in Gene Set |
| --- | --- | --- | --- |
| Wang_adult_alpha.genes | 27364731 | Wang_YJ_et_al_2016_Adult_Alpha_Cell.genes | 212 |
| Wang_adult_beta.genes | 27364731 | Wang_YJ_et_al_2016_Adult_Beta_Cell.genes | 376 |
| Muraro_Alpha.genes | 27693023 | Muraro_MJ_et_al_2016_Alpha_Cell.genes | 572 |
| Muraro_beta.genes | 27693023 | Muraro_MJ_et_al_2016_Beta_Cell.genes | 953 |
| Muraro_Delta.genes | 27693023 | Muraro_MJ_et_al_2016_Delta_Cell.genes | 250 |
| Muraro_PP.genes | 27693023 | Muraro_MJ_et_al_2016_PP_Cell.genes | 157 |
| Muraro_duct.genes | 27693023 | Muraro_MJ_et_al_2016_Duct_Cell.genes | 1280 |
| Muraro_Acinar.genes | 27693023 | Muraro_MJ_et_al_2016_Acinar_Cell.genes | 733 |
| Muraro_mesenchyme.genes | 27693023 | Muraro_MJ_et_al_2016_Mesenchyme_Cell.genes | 684 |
| Muraro_endothelial.genes | 27693023 | Muraro_MJ_et_al_2016_Endothelial_Cell.genes | 362 |
| Bramswig_Alpha_Strong.genes | 23434589 | Bramswig_NC_et_al_2013_Alpha_Cell_Strong.gene | 465 |
| Bramswig_Beta_strong.genes | 23434589 | Bramswig_NC_et_al_2013_Beta_Cell_Strong.gene | 451 |
| Bramswig_Exocrine_Strong.genes | 23434589 | Bramswig_NC_et_al_2013_Exocrine_Cell_Strong.gene | 279 |

**Supplementary Table2:** GSEA enrichment scores for all PEEG Set under study. All alpha cell gene set from three different meta data are significant in A-D-M mutants and highlighted as red.

| NAME | SIZE | ES | NES | NOM p-val | FDR q-val |
| --- | --- | --- | --- | --- | --- |
| **BRAMSWIG_ALPHA_STRONG.GENES** | **446.000** | **0.674** | **1.620** | **<0.001** | **0.025** |
| **MURARO_ALPHA.GENES** | **555.000** | **0.587** | **1.583** | **0.009** | **0.023** |
| **WANG_ADULT_ALPHA.GENES** | **210.000** | **0.693** | **1.573** | **0.004** | **0.021** |
| MURARO_ACINAR.GENES | 727.000 | 0.401 | 1.134 | 0.291 | 0.646 |
| MURARO_DUCT.GENES | 1269.000 | 0.374 | 0.998 | 0.464 | 0.860 |
| MURARO_DELTA.GENES | 243.000 | 0.364 | 0.984 | 0.468 | 0.755 |
| MURARO_PP.GENES | 155.000 | 0.346 | 0.924 | 0.588 | 0.789 |
| BRAMSWIG_EXOCRINE_STRONG.GENES | 277.000 | 0.341 | 0.802 | 0.749 | 0.986 |
| MURARO_BETA.GENES | 930.000 | 0.274 | 0.738 | 0.833 | 1.000 |
| MURARO_ENDOTHELIAL.GENES | 361.000 | 0.317 | 0.662 | 0.845 | 1.000 |
| MURARO_MESENCHYME.GENES | 676.000 | 0.264 | 0.598 | 0.935 | 0.985 |

**Supplementary Table3:** Bramswig et al., expression for HNF1A in normal alpha and beta cells and motif TFs analysis on alpha specific genes. Table a) shows HNF1A gene expression in three alpha and three beta FACs sorted cells. HNF1A gene expression is significant (pval< 0.008) in alpha cells. Bramswig et al., 465 strong alpha specific genes were used to find motif TFs enriched for these genes using online version of GSEA (C3 TFs motif database). Table b) shows two HNF1 TFs motifs are significant for alpha 465 genes.

| Gene name | 53_alpha | 68_alpha | 70_alpha | 53_beta | 68_beta | 70_beta | P-val |
| --- | --- | --- | --- | --- | --- | --- | --- |
| **HNF1A expression** | 1.429458333 | 1.458808333 | 1.58975 | 0.981316667 | 1.18395 | 0.995770833 | 0.007123818 |

**Supplementary Table 3:** Bramswig et al., expression for HNF1A in normal alpha and beta cells and motif TFs analysis on alpha specific genes.
| Motif TFs | p-value | FDR q-value |
| --- | --- | --- |
| **HNF1_01** | 7.33E-06 | 7.64E-05 |
| **RGTTAMWNATT_HNF1_01** | 2.80E-14 | 1.91E-12 |

